# Oscillatory correlates of threat imminence during virtual navigation

**DOI:** 10.1101/2023.09.18.558288

**Authors:** Galit Karpov, Mei-Heng Lin, Drew B. Headley, Travis E. Baker

## Abstract

The Predator Imminence Theory proposes that defensive behaviors depend on the proximity of a threat. While the neural mechanisms underlying this proposal have been well studied in animal models, it remains poorly understood in humans. To address this issue, we recorded EEG from twenty-four (15 female) young adults engaged in a first-person virtual reality Risk-Reward Interaction task. On each trial, participants were placed in a virtual room and then presented with either a threat or reward conditioned stimulus (CS) in the same room location (proximal) or different room location (distal). At a behavioral level, all participants learned to avoid the threat-CS, with most using the optimal behavior to actively avoid the proximal threat-CS (88% accuracy) and passively avoid the distal threat-CS (69% accuracy). By contrast, participants learned to actively approach the distal reward-CS (82% accuracy) and to remain still (passive) to the proximal reward-CS (72% accuracy). At an electrophysiological level, we observed a general increase in theta power (4-8 Hz) over right posterior channel P8 across all CS conditions, with the proximal threat-CS evoking the largest theta response. By contrast, distal CS cues induced two bursts of gamma (30-60 Hz) power over midline-parietal channel Pz (approx. 200 msec post-cue) and right frontal channel Fp2 (approx. 300 msec post-cue). Interestingly, while both bursts were sensitive to distal-CS cues, the first burst of gamma power was sensitive to the distal threat-CS requiring a passive response, and the second burst at channel Fp2 was sensitive to the distal reward-CS requiring an active response. Together, these findings demonstrate that oscillatory processes differentiate between the spatial proximity information during threat and reward encoding, likely optimizing the selection of the appropriate behavioral response.

## Introduction

The ability to identify threats and select an optimal defensive response (e.g., freeze, fight, or flee) is crucial for survival, and has been extensively studied across species using associative learning paradigms (Likhtik & Paz, 2015; Trenado et al., 2018). For example, in Pavlovian fear conditioning tasks a conditioned stimulus [CS] is paired with an aversive stimulus (e.g. aversive tone, foot shock, or odor), and over the course of learning the animal begins to exhibit defensive responses (e.g. freezing or fleeing) when exposed to the CS alone. Decades of electrophysiological recordings from rodents, as well as recent human neuroimaging studies (e.g., functional magnetic resonance imaging, fMRI), have established that the amygdala, medial temporal lobe (e.g. hippocampus, entorhinal cortex, parahippocampal cortex), and prefrontal cortex (e.g. cingulate cortex, orbital frontal cortex, medial prefrontal cortex) are essential components of the circuitry that mediate fear conditioning (LeDoux, 2003; Maren, 2001; Trenado et al., 2018). Moreover, local field potential recordings in animals and non-invasive M/EEG recordings in humans indicate that theta (4-8 Hz) and gamma (30-90 Hz) oscillations provide a mechanism for the fine-tuned organization of the neural circuitry that are engaged during classical fear conditioning, particularly during the coordinated interaction between the prefrontal cortex, medial temporal cortex, and amygdala (Trenado et al., 2018).

Although the neural circuits underlying fear conditioning are increasingly understood, the simplicity of conventional fear conditioning paradigms lowers the ecological validity when compared to real life encounters. For example, in a real-world setting the learning system has the ability to integrate contextual spatial information (i.e., is the threat close or far away) to determine threat imminence, which in turn allow the organism to match the most effective defensive behavior to the current situation (e.g. freeze vs escape). Further, across species it has been shown that behaviors and neural mechanisms differ when a threat is unavoidable, as seen in classical Pavlovian fear conditioning, compared to when a threat is avoidable, as seen in active avoidance paradigms (Boeke et al., 2017; Löw et al., 2015; Moscarello & LeDoux, 2014; Wendt et al., 2017). Accordingly, a central claim of the Predatory Imminence Theory is that defensive behavior changes with increasing threat proximity (Fanselow & Lester, 1988), such that animals are more likely to exhibit freezing or passive avoidance behaviors when threats are distal, but may switch to fighting or fleeing behavior as threats become more proximal (Blanchard & Blanchard, 1988; Deakin & Graeff, 1991; Fanselow & Lester, 1988). Further, in vivo recordings from freely moving rodents indicate distal-threat defensive behaviors rely more on prefrontal regions, whereas proximal-threat defensive behaviors rely more on subcortical regions such as the midbrain (Fanselow & Lester, 1988; Mobbs et al., 2020). While the neural mechanisms underlying distal/proximal threat behaviors have been based mainly on research in freely moving animals, these mechanisms remain poorly understood in humans.

This apparent lack of knowledge is likely due to the limited options for human neuroimaging paradigms to investigate how threat imminence in the ‘real-world’ influences behavioral and neural response patterns. Only recently has the development of novel neuroimaging and virtual 3D design techniques made threat imminence amenable for investigation in humans (Gold et al., 2015; Perkins et al., 2011; Qi et al., 2018; Rodrigues et al., 2018). Further, fMRI studies have revealed that distal-threats encountered in a virtual environment involve the prefrontal cortex (e.g., posterior cingulate cortex) and hippocampal formation, while proximal threats were shown to engage the periaqueductal gray area and midcingulate cortex (Mobbs et al., 2009; Mobbs et al., 2007; Mobbs et al., 2010; Qi et al., 2018). While these studies provide initial evidence of the neural correlates of motivational processes during virtual tasks, they remain limited. To our knowledge, EEG has yet to be utilized to investigate threat imminence (distal vs proximal) in humans. In principle, an EEG experiment could provide critical temporal and spectral information in the context of threat imminence. Given that EEG studies have already reported differential oscillatory activity (gamma and theta) in relation to simple fear conditioning tasks (Trenado et al., 2018), as well as spatial navigation tasks (Baker & Holroyd, 2013; Lin et al., 2022), we suggest that both gamma and theta activity could be appropriate candidates to investigate threat imminence in humans navigating a virtual environment. Thus, to extend previous work, we examined oscillatory activity in humans engaged in a virtual version of the Risk-Reward Interaction task (RRI; (Kyriazi et al., 2018)^1^), in which participants are required to learn the appropriate instrumental behavior (active vs passive responses) according to the context of the CS presented in a virtual room (i.e., valence [threat vs reward] and imminence [distal vs proximal]). Building on the Predatory Imminence Theory, the recording of EEG during the virtual RRI task provides a unique opportunity to examine conditioning-induced alterations in neural activity related to both the valence and imminence of the CS.

## Methods

### Participants

Twenty-four young adults (23 right-handed; Laterality Quotient M = 81.99, SD = 28.43, 9 male and 15 female, aged 18–26 years old [M = 20.87, SD = 2.03]) participated in the study. While twenty-six subjects were initially collected, two participants were removed from the analysis due to excessive artifacts (over 20% of the data) and failing to learn to avoid the threat-CS. Participants were recruited from Rutgers University – Newark Department of Psychology subject pool using the SONA system. Each subject received course credit for their participation and a monetary bonus based on task performance. Participants were screened for current or previous neurological symptoms as well as history of neurological injuries (e.g., head trauma with loss of consciousness longer than 5 minutes) and asked to self-report any psychiatric diagnoses. Participants then completed the Edinburgh Handedness Inventory (Oldfield, 1971). After the experiment, participants filled out the Reinforcement Sensitivity Theory of Personality Questionnaire (Corr & Cooper, 2016), the Behavioral Inhibition System/Behavioral Activation System (BIS/BAS) Scale (Carver & White, 1994), and the CES-D (Radloff, 1977), the results of which will be reported elsewhere.

This study was approved by the Institutional Review Board of Rutgers University and all experiments were performed in accordance with relevant guidelines and regulations. The study adhered to the principles expressed in the 1964 Declaration of Helsinki. Informed consent was obtained from all participants.

### Procedure

The virtual desktop RRI task was constructed using Home design 3D (Expert Software Inc., Coral Gables, FL) and implemented using E-Prime experiment control software (Psychological Software Tools, Pittsburgh, PA). In keeping with the rodent design of the RRI (Kyriazi et al., 2018), this virtual version consisted of a linear brick room (Figure. 1A and 1B) with an open ceiling exposed to a cloudy blue sky. Different brick textures were given to the side and front facing walls. To give participants a first-person video game-like experience, still images were taken from cardinal points inside the 3-dimensional linear room (left, center, and right location). The start image was viewed from the middle of each start location and looking towards to the front wall. All stimuli were presented on a 24-inch monitor (60 Hz refresh rate, 1920 x 1080 resolution), placed approximately 60-70cm from the subjects’ eyes (See supplementary video for an example of the task).

**Figure 1.**
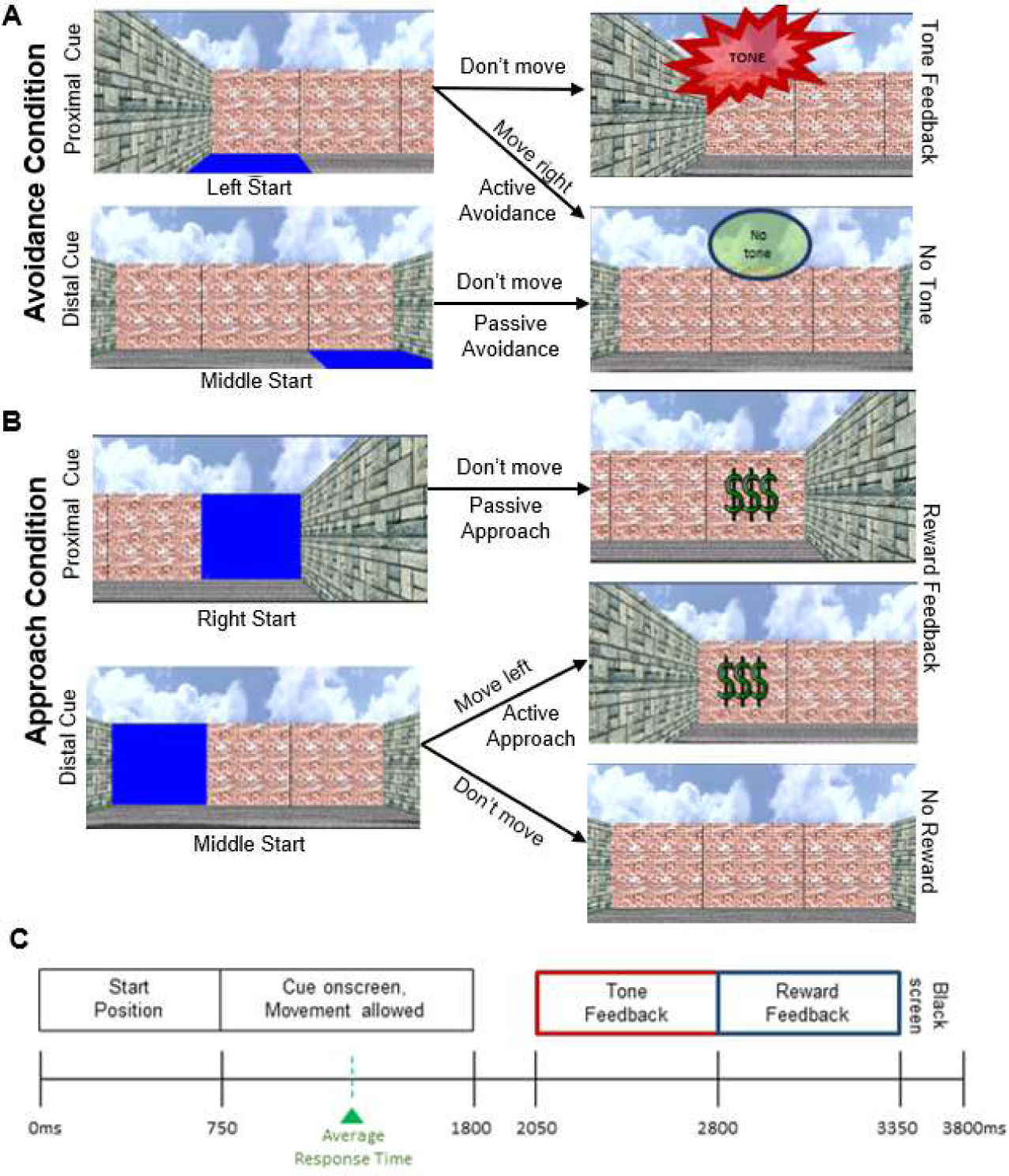
Virtual Risk-Reward Interaction task. **A.** Avoidance condition. Example trial for the Proximal (top panel) and Distal (bottom panel) threat conditions. When the proximal threat-CS was presented, participants learned to move away from the current sector to avoid the aversive tone (active avoidance, bottom right panel). For the distal threat-CS condition, participants learned to remain in place to avoid the aversive tone (passive avoidance, top right panel). **B**. Approach condition. Example trial for the Proximal (top panel) and Distal (bottom panel) reward conditions. When the proximal reward-CS was presented, participants learned to remain in place to receive the reward (5 cents) (passive approach, top right panel). For the distal reward-CS condition, participants learned to approach the reward location to receive the reward (active approach, bottom panel). **C**. Trial timeline for a single trial. Average response time during active conditions was approx. 525msec after CS onset.

Participants were seated comfortably in a sound attenuated chamber under dim lighting and asked to position their right hand so that the fingertips of their index, middle, and ring fingers rested on a mechanical gaming keyboard placed in front of them. Each finger (button) corresponded to a room location. The right index finger moved the participant to the left room location, the right middle finger moved the participants to the center room location, and the right ring finger moved them to the right room location. If they pressed the button corresponding to the location they started in, no movement in the room occurred but the response was recorded as an “active response”. Participants received both written and verbal task instructions explaining the spatial dimensions of the virtual room, and the two response options during CS onset. Importantly, participants were told they had to learn the task through trial-and-error learning, but no detail were provided about the valence dimension of the CS (reward or threat) or the associated optimal response (approach or avoidance, active or passive response). In addition, participants were told that by not pressing any button, they would remain in the current location. Participants were asked to maintain correct posture and minimize muscle movements. (See Supplement S1 for transcript of task instructions).

On each trial, subjects were pseudo-randomly placed in one of the three start locations (left, center, right). After 750 msec, one of two CS would appear in one of the three start locations (Figure. 1). For the threat-CS (Avoidance condition), a section of the floor panel would turn blue (Figure 1A), and for the reward-CS (Approach condition), a section of the wall panel would turn blue (Figure 1B). Participants had to learn that ending in the same section as the reward-CS resulted in a monetary reward ($$$, 5 cents), while ending in the same section as the threat-CS resulted in an aversive buzzard tone (65 decibels) played on a speaker placed 40 cm behind the participant (See supplementary video). During the presentation of the CS (duration: 1050msec), participants were able to make an active or passive response. The theoretical optimal response to the valence (threat vs. reward) and spatial location (proximal vs. distal) of the CS are as follows: For the threat-CS condition, if the CS was presented in the same location as the participant (a proximal threat), the optimal response was to actively move to a different room location to avoid the tone (Active-Avoidance). If the CS was presented in a different location (a distal threat), the optimal response would be to remain in their current location without responding (Passive-Avoidance). An active response to avoid the tone (moving to the other safe location or by pressing a button to stay in place) was also possible, and these responses were recorded and included in the analysis. After 250 msec following the offset of the threat-CS, participants remaining in the same location would receive an aversive tone for 1000 msec. Participants who actively moved to another location would avoid the tone. To note, subjects’ responses were restricted to their first button press.

For the reward-CS condition (Figure 1B), if the CS was presented in the same location (a proximal reward), the optimal response was to remain passively in the same place (Passive-Approach). Any active responses in this condition would result in reward omission (Figure 1B), including pressing a button to remain in the same location. If the CS was presented in a different room location (a distal-reward), participants were required to move to the CS location (Active-Approach) to receive the reward ($$$ - 5 cents, duration 1000 msec). Failing to do so resulted in the omission of the reward. Feedback was followed by a blank screen for 250 msec, and the next trial would begin. For any active response, only the first button press was recorded and changed room locations. Any subsequent responses on the same trial had no impact on the outcome.

In total, the task contained four conditions: Active-Avoidance, Passive-Avoidance, Active-Approach, and Passive-Approach, and was comprised of four blocks (120 trials per block) separated by self-timed rest-breaks. To clarify our conditions regarding CS proximity, the Passive-Avoidance condition reflected a distal threat-CS, the Active-Avoidance condition reflected a proximal threat-CS, the Passive-Approach condition reflected a proximal reward-CS, and the Active-Approach condition reflected a distal reward-CS. Following the experiment, participants received their total bonus (up to $12.00 US dollars) and were administered questionnaires.

### Data acquisition and EEG processing

The EEG was recorded from a montage of 32 active electrodes in accordance to the extended international 10–20 system (Jasper, 1958). Signals were amplified by low-noise electrode differential amplifiers with a frequency response of DC 0.017–67.5 Hz (90 dB octave roll off) and digitized at a rate of 1000 samples per second. Digitized signals were recorded to disk using Brain Vision Recorder software (Brain Products GmbH, Munich). Impedance was kept below 20 kΩ. Two electrodes were placed on the left and right mastoids. The EEG was recorded using the average reference, and Fpz was selected for the ground. The electroocculogram (EOG) was recorded for ocular correction. For pre-COVID participants, the horizontal EOG was recorded from the external canthi of both eyes, and vertical EOG was recorded from the suborbit of the right eye and electrode channel Fp2. During COVID, the horizontal EOG was recorded from electrodes F9 and F10, while vertical EOG was recorded from the supraorbit of the right eye due to masks and to minimize interaction around the face.

Post-processing and data visualization were performed using Brain Vision Analyzer 2 software (Brain Products GmbH, Munich). Muscle artifacts and non-task data (e.g. breaks) were manually removed using the raw data inspector module. The digitized signals were then filtered through a pass-band of 1-60 Hz, order of 4, with a notch filter at 60Hz. A horizontal eye channel (hEOG) was created by subtracting the right from left horizontal EOG electrodes. For pre-COVID data, a vertical eye channel (vEOG) was created by subtracting the vertical EOG electrode from Fp2. For data collected during COVID, no new vertical eye channels were created and the existing vEOG was used. These eye channels were used for the automated ocular ICA algorithm. ICA parameters included a Mean Slope Algorithm, with vEOG and hEOG channels referenced to a common reference using continuous data and Extended Infomax ICA. The convergence bound was set to 1E-07, and number of ICA steps to 512. ICA components were found using Sum of Squared Correlations with vEOG and hEOG, with 30% of the variance captured by the component deleted. After correcting ocular artifacts, data was re-referenced to the linked mastoids, and then segmented −2500 to 2500 msec centered on the CS. Baseline correction was then applied to each segment using a −200 to 0 msec before CS onset. Muscular and other artifacts were automatically removed using a ±150 μV level threshold within a 100 msec interval and a ±35 μV step threshold as rejection criteria. No more than 5% of data was rejected from any channel or condition from any subjects. Channels where more than 5% of data was rejected were interpolated. Lastly, data were segmented into each of the four conditions for the time-frequency decomposition: Passive Approach, Active Approach, Passive Avoidance, Active Avoidance.

### Time-Frequency Processing

The segmented EEG data was convolved with a complex 7-cycle Morlet wavelet using the Wavelet Toolbox (2006) in MATLAB (2019b) as part of custom-written MATLAB code. Frequencies in the 1-60 Hz range were convolved, going through each frequency in linear steps of one. Instantaneous power (squared amplitude) at each time point was computed for each trial and then averaged within each subject across conditions and blocks. The relative change in the power for each condition was determined by averaging the baseline activity (−200 to −100 msec pre-stimulus) across time for each frequency and then subtracting the average from each data point following stimulus presentation for the corresponding frequency. This value was then divided by the baseline activity to normalize the change of power to the baseline activity [Equation: normalized power = (raw power-baseline)/baseline]. For each subject, the total frequency power was calculated as the average value of time-frequency power across single trials and the evoked power was determined directly from the averaged ERPs (Behroozmand et al., 2015; Hajihosseini & Holroyd, 2013).

Based on previous animal (e.g., local field potentials) and human (e.g., M/EEG) findings during fear conditioning tasks (Trenado et al., 2018), we focused our analysis on theta (4-8 Hz) and gamma (30-60 Hz) activity following the onset of the CS. For each frequency band, we first found the scalp location where frequency power was maximal by averaging the EEG data across all conditions and identified the peak power within a 400 msec time window after CS onset (Figure 2, Table S1). We selected the channel location where evoked theta power (P8) and total gamma power (Fp2 and Pz) was largest for the subsequent analysis (Figure 2, Table S1). Spectral power was averaged within a 50 msec window around the peak of theta (236 msec - P8), and the peak of two gamma bursts (approx. 300 msec - Fp2 | 210 msec – Pz). Because a Shapiro-Wilk test indicated a positive skew across our measures (evoked theta at P8: skew = 3.12, W = 0.70, p < .05, total gamma at Fp2: skew = 3.87, W = 0.58, p < .05, total gamma at Pz: skew = 3.10, W = 0.73, p < .05), the normalized power values were transformed by adding 1 to the power to remove negative values, then the log of those values [Equation: transformed data = ln (normalized power + 1)] were used to run the parametric statistics. This improved the distribution to better meet parametric assumptions.

**Figure 2.**
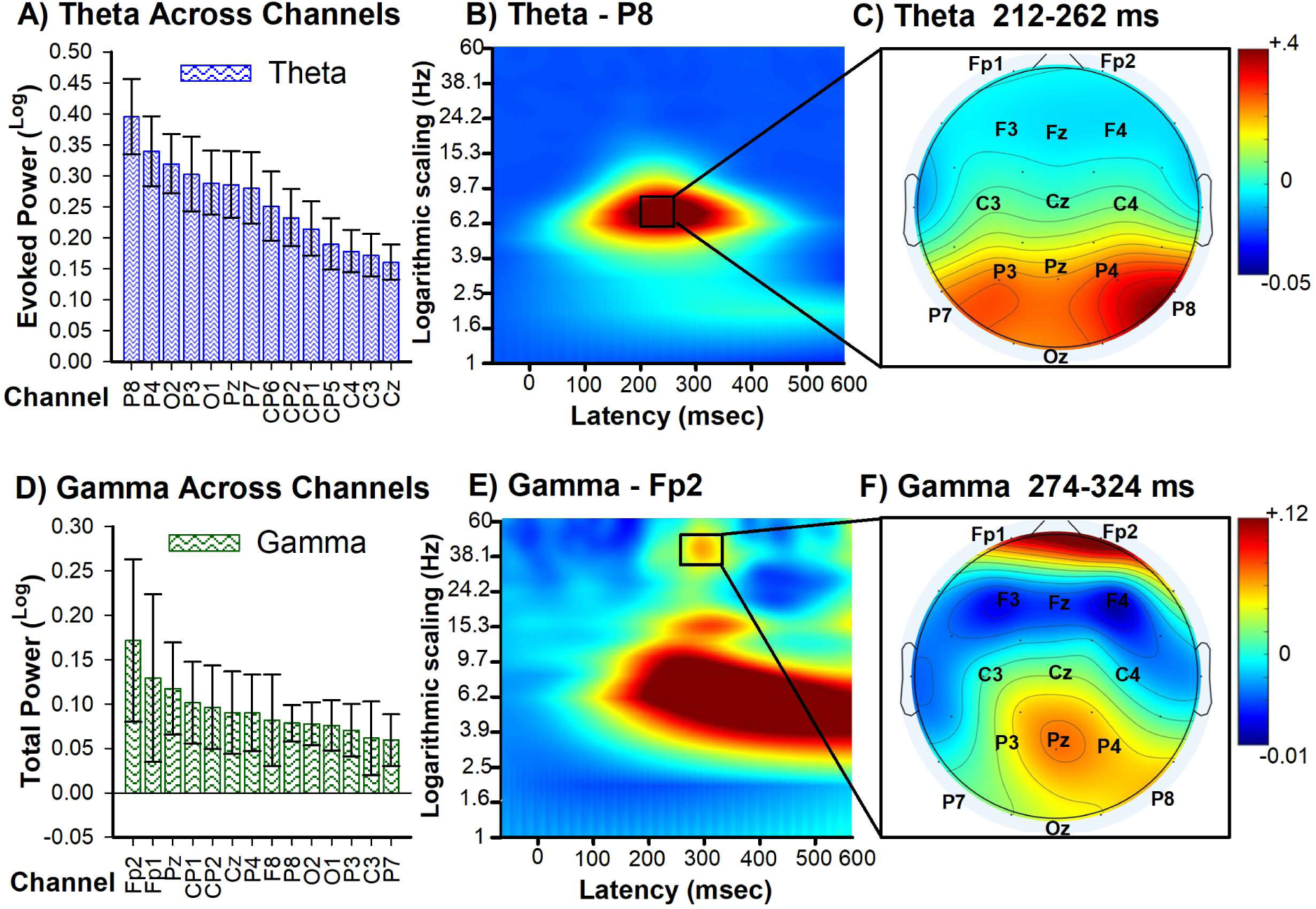
Condition stimulus (CS) processing during virtual navigation. **A**. Bar graph illustrates the peak log power (0–400 msec) for theta (4-8 Hz) activity averaged across all conditions and evaluated at each electrode channel (14 of the 27 channels depicted here) and ordered by size. **B.** Panels indicate changes in power for each frequency band with respect to baseline (− 200 to − 100 msec period prior to CS onset) elicited by the CS cue (averaged across conditions) at channel P8 where theta was strongest. Spectral power is unitless because it is calculated as a proportional increase/decrease relative to baseline. **C.** Topographical maps representing the mean frequency power (212-262 msec) post-CS onset at each channel for theta [4–8 Hz]. **D**. Bar graph illustrates the peak log power (0–400 msec) for gamma (30-60 Hz) activity averaged across all conditions and evaluated at each electrode channel (14 of the 27 channels depicted here) and ordered by size. **E.** Panels indicate changes in power for each frequency band with respect to baseline (− 200 to − 100 msec period prior to CS onset) elicited by the CS cue (averaged across conditions). Data recorded at channel Fp2 where gamma was strongest. **F.** Topographical maps representing the mean frequency power (274-325 msec) post-CS onset at each channel for gamma [30–60 Hz]. Bars indicate the standard error of the mean.

### Statistical Analysis

Statistics were run using the R computing environment (version 4.0.5), with the ‘*rstatix*’ library (version 0.7.0) and SPSS 18.0.1 for Windows (SPSS, Inc., Chicago, IL). Violation of sphericity assumptions were corrected using the Greenhouse-Geisser method. Two-tailed pairwise t-tests were run when a significant ANOVA effect was observed. Type 1 errors were controlled following Benjamini–Hochberg method with a corrected significance level of α = .05. For EEG analysis, all trials were included regardless of correct or incorrect outcomes. Repeated measures multifactorial ANOVAs, type III, were run on the log-transformed power. The 4×2×2 design included Block (Block 1-4), Valence (avoidance [threat] vs. approach [reward]) and theoretical optimal Action (passive vs. active) as within-subject factors. Non-parametric permutations tests were run on the data to ensure that the parametric statistic results were reflective of the normalized power. Methods for the non-parametric statistics can be found in Supplement S2, with supplementary figures S2, S3, S4 showing the clusters and p-values for evoked theta at P8 and total gamma at Pz and Fp2 respectively.

## Results

### Behavioral results

RRI task performance measures included accuracy and reaction time (RT: reported in milliseconds) for overall performance (Approach trials receiving the monetary reward and Avoidance trials without receiving the aversive tone, Figure 3A) and optimal performance (withholding a button response on Passive Approach and Avoidance trials, Figure 3B). In regards to overall RRI task accuracy (Figure 3A and 3C), a repeated measures ANOVA with Block (Block 1-4), Valence (Avoidance, Approach), and Action (Active, Passive) as factors revealed a main effect of Block, F_(3,69)_ = 30.98, p < .001, η_p_^2^ = 0.57, indicating a general increase in performance accuracy following the first block (x̄ = 77%, SE = 3) of trials ̶ Block 2 (x̄ = 90%, SE = 3; p < .001), Block 3 (x̄ = 93%, SE = 2; p < .001), and Block 4 (x̄ = 93%, SE = 3; p < .001). A main effect of Action was also observed, F_(1,23)_ = 26.43 p < .001, η_p_^2^ = 0.54, indicating that participants were more accurate on trials requiring a passive response (x̄ = 91%, SE = 2) relative to an active response (x̄ = 86%, SE = 3; p < .001) (but see optimal performance results). Finally, an interaction was detected between Block and Action, F_(3,69)_ = 5.01, p < .01, η_p_^2^ = 0.17. Post-hoc tests indicated that between Block 1 and Block 2, the Active condition showed a larger accuracy increase (Δx̄ = +17%, SE = 2) compared to the Passive condition (Δx̄ = +10%, SE = 2; p < .01). No other accuracy differences between Actions were observed across Blocks (p > .05). No other main effects or interactions were observed. In regards to RT on Active conditions (Figure 3D and 3E), a repeated measures ANOVA with Block (Block 1-4) and Valence (Avoidance, Approach) as factors revealed a main effect of Block, F_(3,69)_ = 8.13, p < .001, η_p_^2^ = 0.26, indicating that relative to the first two blocks (Block 1: x̄ = 549 msec, SE = 12 | Block 2: x̄ = 532 msec, SE = 12), there was general decrease in RT at Block 3 (x̄ = 514 msec, SE = 13; p < .005 | p < .01) and Block 4 (x̄ = 510 msec, SE = 12; p < .005 | p < .005), respectively. No other main effects or interactions were observed.

**Figure 3.**
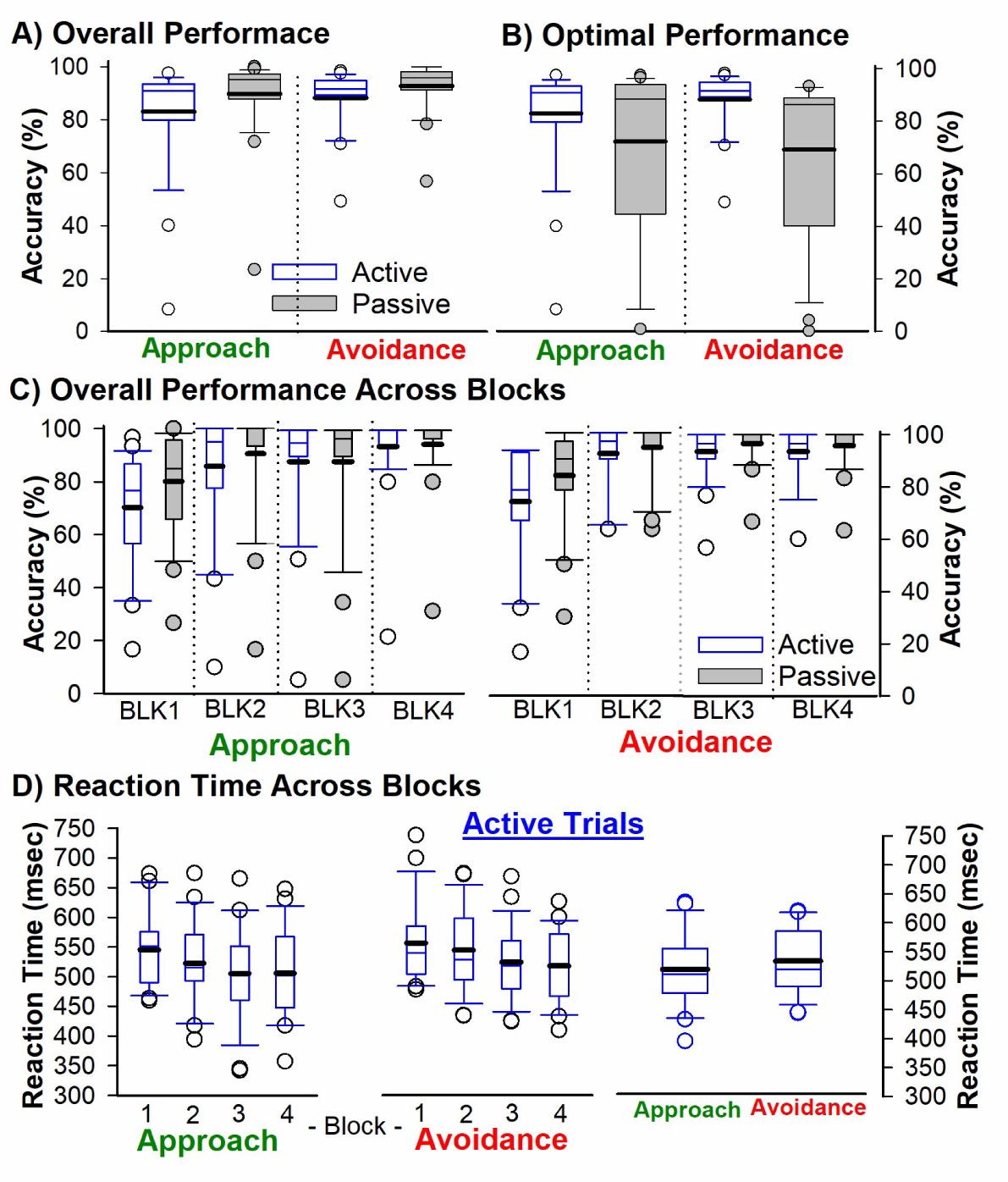
Descriptive Statistics for RRI behavioral performance. **A.** Overall Active (Blue Bars) and Passive (Gray Bars) response accuracy as measured by percentage of Approach trials receiving the monetary reward, and Avoidance trials without receiving the aversive tone. **B**. Percentage of true optimal passive (no button press) and active response in the approach and avoidance condition. **C.** Overall active and passive response accuracy across blocks for the approach (left panel) and avoidance (right panel) conditions. **D**. Reaction time for the overall Active response for the Active Approach (left panel) and Active Avoidance (center panel) conditions across blocks and averaged across blocks (right panel). Box plots represent statistical values. The boundary of the box closest to zero indicates the 25th percentile, a thin line within the box marks the median, the thick line within the box marks the mean, and the boundary of the box farthest from zero indicates the 75^th^ percentile. Whiskers (error bars) above and below the box indicate the 90th and 10th percentiles, and symbols denotes outlying points outside the 10th and 90th percentiles.

While the results above indicate that the majority of participants learned to avoid the threat-CS and approach the reward-CS, not all participants engaged in the theoretical optimal response during the passive conditions (See Figure 3B and Figure S1). For instance, while a response was not required for the passive conditions, approximately 20% (n = 5) of participants (i.e., passively responding on less than 50% of trials) continued to actively respond to the distal threat-CS at Block 4 by either pressing a button to stay in the same location or move to another safe location (Figure S1B, right panel). Further, approximately 16% (n = 4) of participants (i.e., accuracy less than 36%) continued to make a response to the proximal reward-CS at Block 4 (Figure S1B, left panel), even though rewards are omitted on such trials. To examine this further, we conducted a repeated measure ANOVA on optimal performance accuracy with Block and Valence as factors. This analysis revealed a main effect of Block, F_(3,69)_ = 22.34, p < .001, η_p_^2^ = 0.50, indicating that relative to the first block (x̄ = 55%, SE = 6), there was general increase in accuracy at Block 2 (x̄ = 74%, SE = 7; p < .001), Block 3 (x̄ = 76%, SE = 7; p < .001), and Block 4 (x̄ = 79%, SE = 6; p < .001). This result indicates that overall, participants successfully learned the optimal response to distal threat-CS cues, and the optimal response to proximal reward-CS cues. No other main effects nor interactions were observed (p > .05)^2^.

### Theta Power Results

Visual inspection of Figure 2A and 2C shows a clear enhancement of theta power over right posterior channel P8. A repeated measures ANOVA on theta power at channel P8 with Block (Block 1-4), Valence (Avoidance, Approach), and Action (Active, Passive) as factors revealed a main effect of Block, F_(3,69)_ = 6.23, p < .005, η_p_^2^ = 0.21, indicating that relative to the first block, there was a decrease in theta power at Block 3 (p < .01) and Block 4 (p < .005) (Figure 8A). A main effect of Valence was also observed, F_(1,23)_ = 12.30, p < .005, η_p_^2^ = 0.35, indicating a larger increase in theta power for Avoidance trials (x̄ = 0.34, SD = 0.27) relative to Approach trials (x̄ = 0.24, SD = 0.25) (Figure 4A). Next, this analysis revealed a main effect of Action, F_(1,23)_ = 7.63, p < .01, η_p_^2^= 0.25, indicating a larger increase in theta power for the Active condition (x̄ = 0.33, SD = 0.28) relative to the Passive condition (x̄ = 0.25, SD = 0.26). Finally, this analysis revealed an interaction between Valence and Action, F_(1,23)_ = 27.5, p < .001, η_p_^2^= 0.55 (Figure 4C, 7A, and 8A). Post-hoc tests indicated that for the threat-CS conditions, there was a greater increase in theta power for Active Avoidance (x̄ = 0.48, SD = 0.29) relative to Passive Avoidance (x̄ = 0.20, SD = 0.17), p < .001. By contrast, for the reward-CS conditions, Passive Approach displayed a stronger theta response (x̄ = 0.30, SD = 0.32) relative to Active Approach (x̄ = 0.17, SD = 0.14, p < .05). These findings indicate that theta activity was driven by the imminence of the CS since both Active Avoidance and Passive Approach were proximal cues. Furthermore, it is worth noting that the Active Avoidance condition exhibited the largest burst of theta power than any other condition (p < .001), possibly reflecting the salience of the threat-CS and associated action selected (Figure 7A). No other interactions were observed (p > .05).

**Figure 4.**
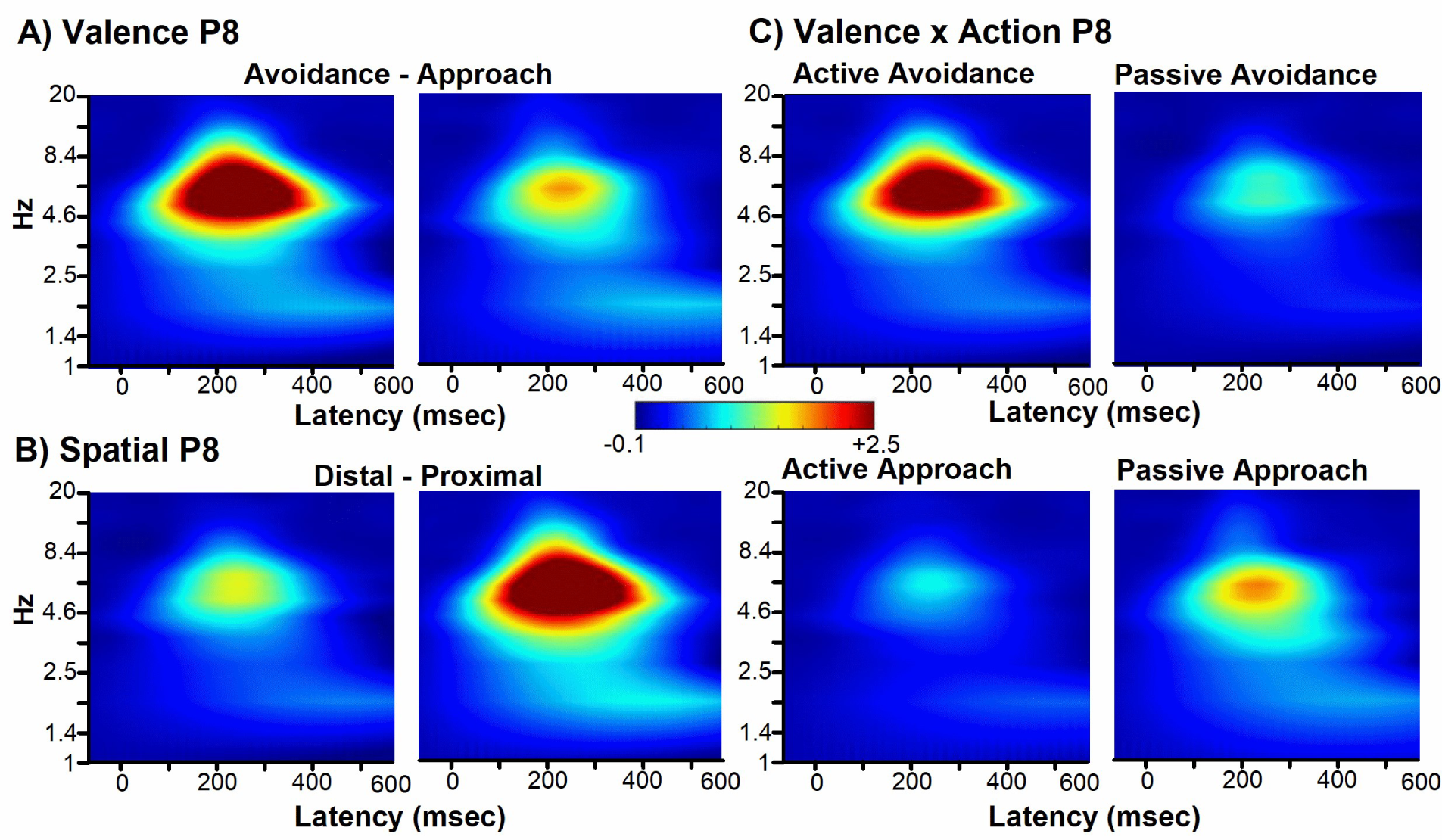
Time–frequency analysis of the EEG data recorded at P8 and time-locked to CS onset. Panel indicates changes in evoked power for each frequency band (1-20 Hz) by condition with respect to baseline. **A)** Panels indicate changes in evoked power associated with Avoidance (left panel) and Approach (right panel) CS conditions. **B)** For illustrative purposes only, panels indicate changes in evoked power associated with Distal (left panel) and Proximal (right panel) CS conditions. **C)** Panels indicate changes in evoked power associated with Active Avoidance (top left), Passive Avoidance (top right), Active Approach (bottom left) and Passive Approach (bottom right) conditions. Spectral power is unitless because it is calculated as a proportional increase/decrease relative to baseline (Hajihosseini & Holroyd, 2013).

### Gamma Power

As shown in Figure 2B and Table S1, two successive bursts of gamma power were observed over posterior midline channel Pz (peaking around 210 msec) and right frontal channel Fp2 (peaking around 300 msec) following CS onset. In regards to the first gamma burst (Figure 5), a repeated measures ANOVA on total gamma power at channel Pz with Block (Block 1-4), Valence (Avoidance, Approach), and Action (Active, Passive) as factors revealed a main effect of Block, F_(3,69)_ = 7.40, p < .001, η_p_^2^ = 0.24, indicating a decrease in gamma power at Block 4 relative to Block 3 (p < .01), Block 2 (p < .01), and Block 1 (p < .001; B–H corrected, p < .025) (Figure 8B). This analysis also revealed a main effect of Valence, F_(1,23)_ = 9.54, p = .005, η_p_^2^ = 0.29, indicating an stronger increase in gamma power for the Avoidance condition (x̄ = 0.095, SD = 0.34) relative to the Approach condition (x̄ = −0.013, SD = 0.24) (Figure 5A). A main effect of Action was also observed, F_(1,23)_ = 4.67, p < 0.05, η_p_^2^ = 0.17, indicating greater power for the Passive condition (x̄ = 0.066, SD = 0.34) relative to the Active (x̄ = 0.016, SD = 0.24) condition. It is worth noting that the Passive Avoidance condition exhibited the strongest gamma burst compared to any other condition (p < 0.05: Figure. 5C, 7B). Furthermore, this analysis revealed an interaction between Valence and Action, F_(1,23)_ = 20.34, p < .001 η_p_^2^ = 0.47. Post-hoc analysis indicated that within the CS-threat condition, Passive Avoidance displayed a stronger gamma burst (x̄ = 0.21, SD = 0.39) relative to the Active Avoidance condition (x̄ = - 0.026, SD = 0.24; p. < .001). By contrast, within the reward-CS condition, Active Approach displayed a larger burst of gamma power (x̄ = 0.057, SD = 0.25) relative to Passive Approach (x̄ = −0.083, SD = 0.21; p. < .001, B–H corrected, p < .0125) (Figure 8B). To note, this interaction suggests that gamma was sensitive to the proximity of the CS since both Passive Avoidance and Active Approach had distal cues. Finally, an interaction between Block and Valence was observed, F_(1,23)_ = 20.34, p < .001 η_p_^2^ = 0.47. Post-hoc tests indicated that there was a general decrease in gamma power across blocks for the Avoidance condition, but not for the Approach condition (Figure 8B). No other interactions were observed (p > .05)

**Figure 5.**
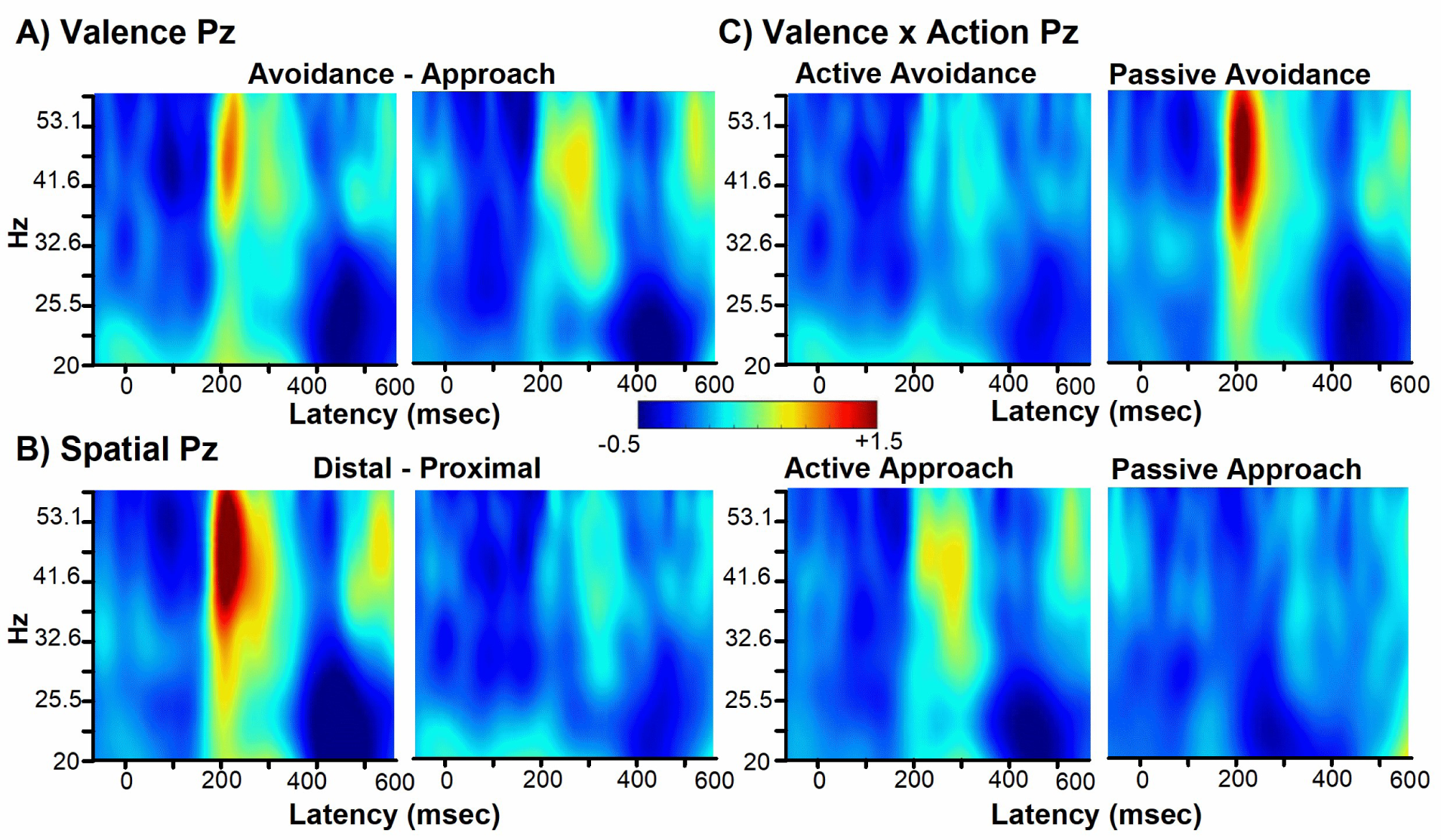
Time–frequency analysis of the EEG data recorded at Pz and time-locked to CS onset. Panel indicates changes in power for each frequency band (20-60 Hz) by condition with respect to baseline. **A)** Panels indicate changes in total power associated with Avoidance (left panel) and Approach (right panel) CS conditions. **B)** For illustrative purposes only, panels indicate changes in total power associated with Distal (left panel) and Proximal (right panel) CS conditions. **C)** Panels indicate changes in total power associated with Active Avoidance (top left), Passive Avoidance (top right), Active Approach (bottom left) and Passive Approach (bottom right) conditions. Spectral power is unitless because it is calculated as a proportional increase/decrease relative to baseline (Hajihosseini & Holroyd, 2013).

In regards to the second burst of gamma at approx. 300 msec over channel Fp2 (Figure 6), a repeated measure ANOVA revealed an interaction between Valence and Action, (F_(1,23)_ = 5.43, p < .05, η_p_^2^ = 0.20 (Figure 6C and 7C). However, unlike the results reported for the earlier posterior gamma response, the comparisons within the CS-reward condition (Active Approach [x̄ = 0.18, SD = 0.45] versus Passive Approach [x̄ = −0.04, SD = 0.18; p = .03]) and threat-CS condition (Passive Avoidance [x̄ = 0.10, SD = 0.35] versus Active Avoidance [x̄ = −0.02, SD = 0.16; p < .04], did not meet statistical threshold when controlling for multiple comparisons (B–H corrected, p < .0125). Instead, follow-up analysis revealed that Active Approach displayed a stronger gamma burst relative to the Active Avoidance condition (p = .009, corrected for multiple comparisons, B– H, p < .0125), and no differences were observed between Passive Approach and Passive Avoidance (p = .10). Thus, while there was an observable difference in right frontal gamma between Distal-cues and Proximal-cues (a posteriori comparison: p < .05, Figure 8C), the interaction reported here possibly reflects the salience of the reward-CS when adaptive actions are required (Figure 7C and 8C). Finally, this analysis also revealed a three-way interaction between Block, Valence, and Action, F_(3,69)_ = 6.62, p < .001, η_p_^2^ = 0.22. Post-hoc tests indicated that for the Reward-CS condition, no differences in gamma power were detected between Active and Passive Approach conditions at Block 1 (p > .05) but was significantly different at Block 2 (p < .005) and Block 3 (p < .05) and Block 4 (p < .05), B–H corrected, p < .025). (Figure 8C). By contrast, this pattern of results was not observed for the threat-CS condition (p > .05). To note, when comparing Active Approach versus Active Avoidance across blocks, all blocks revealed a difference between the two conditions (Block 2: p < .01, Block 3: p < .005, Block 4, p < .05; B–H corrected, p < .037) except Block 1 (p > .05). Together, this interaction suggests that frontal gamma power was particularly sensitive to the Active Approach CS following the initial block of trials and was maintained throughout the experiment. No other main effects or interactions were observed (p > .05).

**Figure 6.**
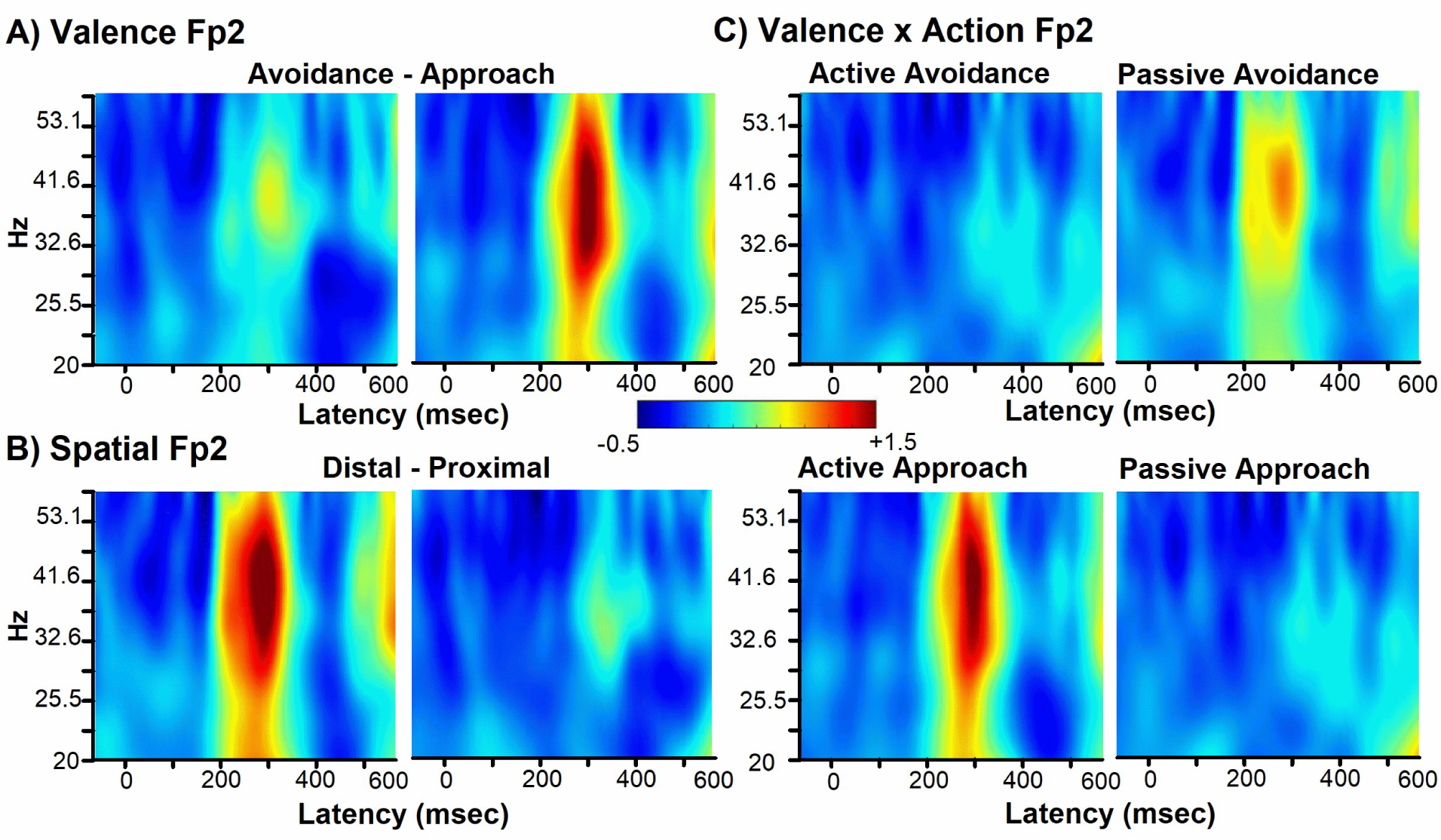
Time–frequency analysis of the EEG recorded at Fp2, associated with CS processing. Panel indicates changes in power for each frequency band (20-60 Hz) by condition with respect to baseline. **A)** Panels indicate changes in total power associated with Avoidance (left panel) and Approach (right panel) conditions. **B)** For illustrative purposes only, panels indicate changes in total power associated with Distal (left panel) and Proximal (right panel) conditions. **C)** Panels indicate changes in total power associated with Active Avoidance (top left), Passive Avoidance (top right), Active Approach (bottom left) and Passive Approach (bottom right) conditions. Spectral power is unitless because it is calculated as a proportional increase/decrease relative to baseline (Hajihosseini & Holroyd, 2013).

**Figure 7.**
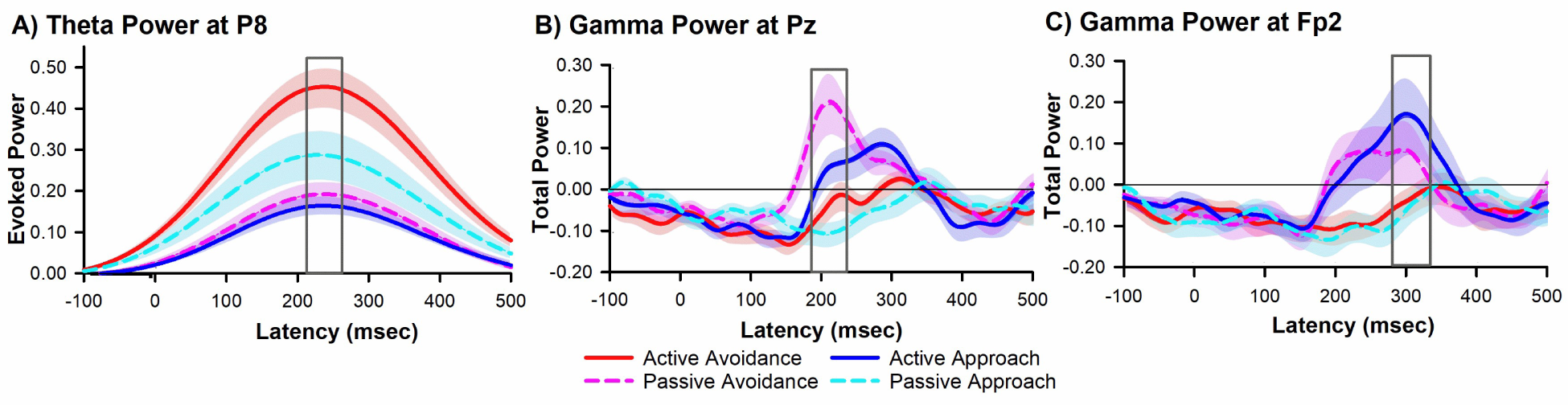
The time course of the change in **A)** evoked theta power recorded at P8, **B)** total gamma power recorded at Pz, and **C)** total gamma power recorder at Fp2 associated Active Avoidance (red solid line), Passive Avoidance (dashed pink line), Active Approach (blue solid line) and Passive Approach (cyan dashed line) conditions. Shaded area denotes the standard error of the mean (SEM) for every datapoint from −100 to 500 msec centered on CS onset.

**Figure 8.**
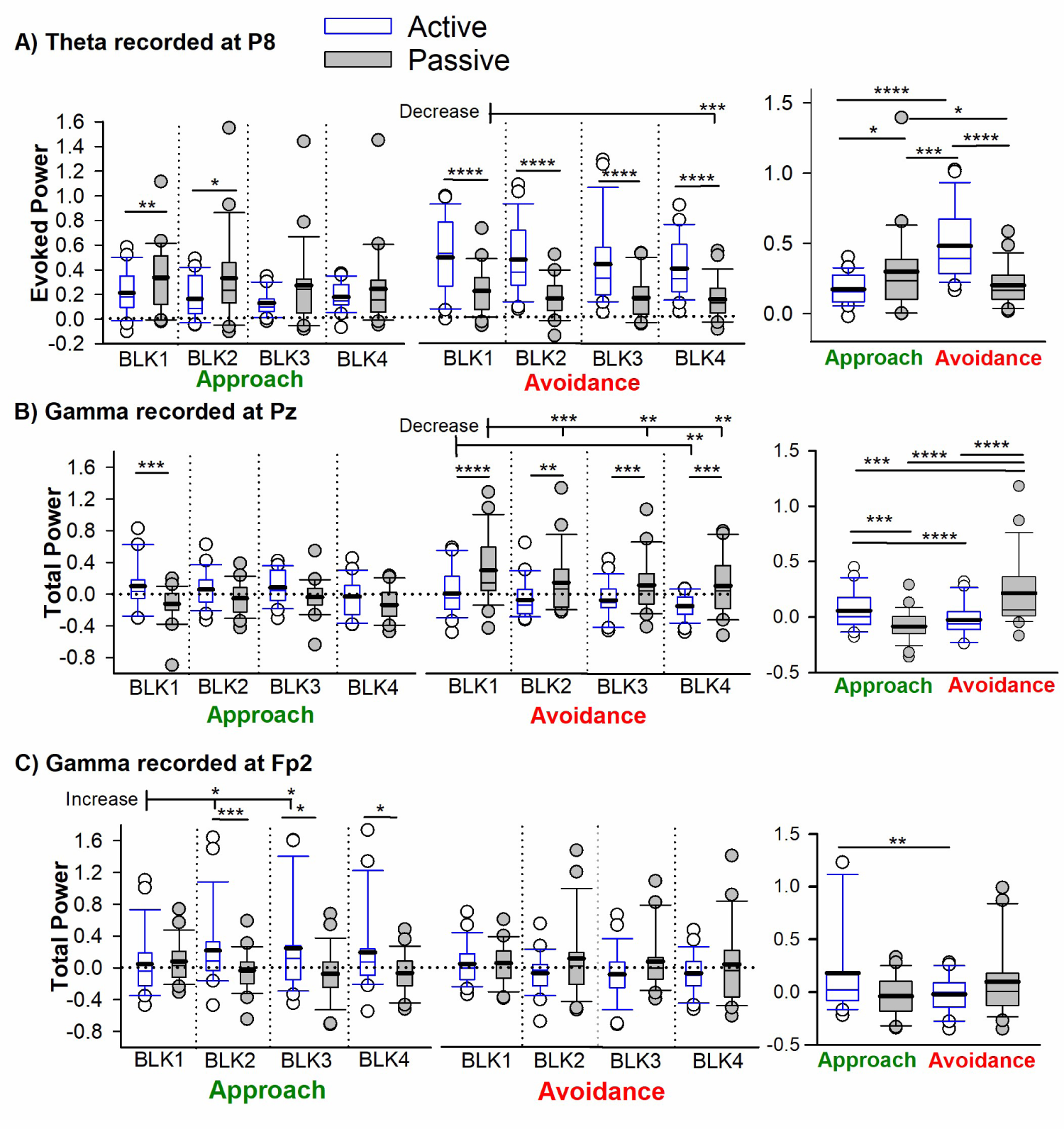
Descriptive Statistics for Frequency power. **A.** Theta power for Active (Blue Bars) and Passive (Gray Bars) trials measured across blocks for Approach (left panel) and Avoidance (central panel) conditions. Right panel denotes theta power for active and passive trials averaged across blocks for the approach and avoidance conditions. **B.** Gamma power recorded at Pz for Active (Blue Bars) and Passive (Gray Bars) trials measured across blocks for Approach (left panel) and Avoidance (central panel) conditions. Right panel denotes gamma power for active and passive trials averaged across blocks for the approach and avoidance conditions. **C.** Gamma power recorded at Fp2 for Active (Blue Bars) and Passive (Gray Bars) trials measured across blocks for Approach (left panel) and Avoidance (central panel) conditions. Right panel denotes gamma power for active and passive trials averaged across blocks for the approach and avoidance conditions. Box plots represent statistical values. The boundary of the box closest to zero indicates the 25th percentile, a thin line within the box marks the median, the thick line within the box marks the mean, and the boundary of the box farthest from zero indicates the 75th percentile. Whiskers (error bars) above and below the box indicate the 90th and 10th percentiles, and symbols denotes outlying points outside the 10th and 90th percentiles. * < .05, ** <.01, *** <.005, and **** <.001: B-H corrected significance level of α = .05.

## Discussion

Predator Imminence Theory is predicated on the idea that defensive actions are determined by threat imminence. Accordingly, animals are more likely to exhibit passive avoidance behaviors when threats are distal, but may switch to fighting or fleeing behavior as threats become more proximal, processes mediated by multiple interacting neural systems (Blanchard & Blanchard, 1988; Deakin & Graeff, 1991; Fanselow & Lester, 1988). Although the neural mechanisms underlying threat imminence are becoming increasingly understood in non-human animals, less is known about this process in humans. In the current study, we investigated whether threat imminence readily modifies electrophysiological response patterns and subsequent actions during an ecologically relevant RRI task. Consistent with the Predator Imminence Theory, our findings demonstrate that the imminence of a threat-related CS differentially modulated electrophysiological response patterns and guided behavioral adaptation in the virtual RRI task.

Foremost, our findings revealed that the presentation of the CS in a virtual environment evoked a burst of EEG oscillations in the theta frequency range (4-8Hz) over right posterior areas of the human scalp. This right posterior theta response decreased over time, consistent with an effect of learning (Takehara-Nishiuchi et al., 2011). Strikingly, the strongest theta response was elicited by the threat-related CS cue encountered in the same room location as the participant (i.e., proximal threat), in which participants quickly learned to actively move to another room location to avoid the impending aversive tone. Further, the proximal reward-CS, which required the subjects to learn to remain in the same location (passive response), also evoked a relatively large right posterior theta response. Together, these findings provide evidence for the existence of a theta mechanism as sensitive to threatening events requiring a defensive action, as well as the spatial proximity of salient events with respect to an egocentric frame of reference. In regard to the threat-sensitivity explanation, power increases in theta oscillations at posterior and latero-frontal channels have been associated with fear following conditioned and unconditioned stimuli in other work, and has been shown to play a role in generating and maintaining fear states, as well as retrieval of fear-evoking stimuli (Chen et al., 2021; Mueller et al., 2014; Pape et al., 2005; Sperl et al., 2018; Trenado et al., 2018). More so, converging evidence from animal and human studies indicate that theta oscillations may provide a mechanism for the coordination of neural circuits engaged in memory formation and recall of fearful events, particularly by emphasizing the interaction between cortical and subcortical regions through theta synchrony (Hyman et al., 2011; Marques et al., 2022; Padilla-Coreano et al., 2019). Medial temporal theta has been shown to be especially important in active avoidance paradigms. For example, hippocampal theta response is larger to a CS when the outcome can be avoided, compared a CS that predicts an unavoidable outcome (Marques et al., 2022). Hippocampal theta is also able to modulate avoidance behaviors, where theta stimulation increases avoidance behaviors (Padilla-Coreano et al., 2019). While the hippocampus cannot be recorded via EEG, it shows strong functional connectivity with cortical regions. This includes the parahippocampal cortex, which is involved in escaping proximal but not distal threats (Qi et al., 2018 – supplementary material).

Regarding the spatial explanation, the latency (approx. 190 - 240 msec) and scalp distribution (P8) of this theta response to proximal CS cues (and all CS cues in general) bear a very close resemblance to a previously observed right posterior theta (RPT) response elicited by salient events encountered during virtual navigation (Baker & Holroyd, 2013; Lin et al., 2022). Across a series of EEG studies, we previously demonstrated that salient events (e.g., rewards, reward-predicting cues, and landmarks) presented during virtual navigation elicit a burst of theta over right posterior electrode locations (P8 and PO8) around 150-250 msec (Baker & Holroyd, 2013; Lin et al., 2022). Furthermore, the spectral dynamics of RPT are highly sensitive to the participant’s specific egocentric orientation in relation to reward-related cues, even in freely moving humans navigating an immersive virtual T-maze (Lin et al., 2022). EEG source localization analysis, fMRI data, and combined EEG-fMRI data point to the right medial temporal cortex, particularly the right parahippocampal cortex, as the most likely source of RPT (Baker & Holroyd, 2013; Baker et al., 2015; Gueth et al., 2019), results that converge with a vast literature indicating that parahippocampal theta activity contributes to spatial learning and memory (Baumann & Mattingley, 2021; Epstein et al., 2017; Weniger et al., 2010). In line with previous animal and computational work, we have argued that RPT is generated by a partial phase reset of the ongoing theta rhythm in the parahippocampal region, and reflects a neural mechanism for encoding contextual and spatial information about salient events such as finding rewards (Baker & Holroyd, 2009; Baker & Holroyd, 2013; Baker et al., 2015).

By extension, this proposal dovetails with previous conceptions about the role of parahippocampal region (i.e., entorhinal cortex, perirhinal cortex, and parahippocampal cortex) in fear conditioning (Alvarez et al., 2008; Baldi & Bucherelli, 2014; East et al., 2022; Greco & Liberzon, 2016; Maren et al., 2013). In particular, while much attention in the field has focused on the contributions of subcortical brain regions to fear conditioning, especially the amygdala and hippocampus, there is growing evidence for cortical involvement – including the parahippocampal cortex – in encoding context during fear conditioning and retention (Maren et al., 2013). Current thinking holds that the parahippocampal region can be considered a gateway that passes contextual information about specific events to and from the hippocampus and that stores memories related to this information (M. Li et al., 2016; Rudy, 2009). Further, it has been suggested that the amygdala is well positioned to evaluate context-dependent threats because of its recurrent connectivity with the parahippocampal-hippocampal system. Accordingly, the parahippocampal cortex, in conjunction with amygdala and hippocampus, mediates the strong connection between contextual processing and threat, facilitating threat encoding and expectations of our environment (Maren et al., 2013; Vishnu et al., 2012). While both amygdala and hippocampal threat-related activities are undetectable at the scalp due to their closed electromagnetic fields, it is nevertheless the case that activity of neighboring areas of medial temporal cortex, particularly the parahippocampal cortex, are amenable to investigation with EEG during threat-related processes. And while not given much attention, the right parahippocampal gyrus has been found to be activated during proximal as opposed to distal threats (Qi et al., 2018; Wendt et al., 2017 – supplementary material for both). Thus, given the morphological and functional similarities between the previously reported spatial RPT response and the right posterior theta response reported here suggests that the two oscillatory components are manifestations of the same underlying phenomenon - the encoding of contextual information about salient events during virtual navigation. Because the parahippocampal region produces a theta rhythm during event encoding, and previous human neuroimaging studies shows greater activation in this region during the processing of threat-related stimuli (Meyer et al., 2019; Qi et al., 2018; Wendt et al., 2017), we tentatively suggest that encountering a proximal threat in the virtual room evoked activity in the right parahippocampal gyrus that in turn produced the right posterior theta response. Taken together, these results provide preliminary evidence for the proposition that RPT reflects a parahippocampal mechanism for encoding contextual and spatial information about threating events for the purpose of learning the most appropriate defensive response. Although admittedly speculative, we hope that this suggestion will motivate future neuroimaging work, particularly focusing on the differential role of RPT in facilitating various defensive strategies.

Regarding gamma, our study revealed two successive bursts of gamma activity (30-60Hz) that reflected a sensitivity to the proximity of the CS specific to distal cues. The earlier posterior-midline gamma burst (approx. 210 msec post-CS onset) was sensitive to both Passive Avoidance and Active Approach conditions, but stronger for Passive Avoidance. Given that these distal cues were linked to specific behavioral contingencies for each condition, this interaction suggests that the observed gamma activity was specific to the neural representations of distal cues in relation to the participants egocentric frame of reference, as well as the motivational significance of the cues. In line with this proposal, a multitude of studies have indicated that parietal gamma activity reflects visuospatial processing during navigation (White et al., 2012), the maintenance and precision of working memory (Morgan et al., 2011; Thompson et al., 2021), re-directing attention to task-relevant stimuli (Gruber et al., 1999), increased emotion processing of cues (Kang et al., 2012), and visuomotor encoding mechanisms that determine the planned direction of the movement towards a goal (Jurrian Van Der et al., 2008; Urai & Donner, 2022). This accumulating body of evidence indicates that gamma serves to bind distributed neural systems (e.g., perceptual systems, memory systems, motivational/reward systems, and motor systems) into functional networks for the purpose of producing learned behaviors (Headley & Pare, 2013; Tallon-Baudry & Bertrand, 1999). However, given the number of different processes and diverse classes of stimuli, tasks, and environments related to the gamma signal, efforts to define a single function for the gamma activity observed in the current study is challenging. Although our gamma findings confirm theoretical predictions that proximal and distal cues are processed differently in the brain, these gamma findings are entirely novel and will need to be explored in future experiments. Nevertheless, on the basis of the current evidence, we tentatively suggest that the neural system generating the parietal gamma activity processed the CS-related information relative to an egocentric frame of reference (distal) and the encoding of the behavioral context (avoid or approach).

By contrast, while the later right-frontal gamma burst (approx. 300 msec post-CS onset) also displayed a sensitivity to distal cues, this response was most pronounced for the Active Approach condition. This possibly reflects the salience of the reward-CS when actions are required. Lateralized frontal activity has been reported in the context of virtual motivated navigation. Rodrigues and colleagues (2018) recorded EEG during a chase-and-capture videogame tasks and found a stronger right frontal activation when encountering a conditioned stimulus indicating that they had to escape a monster to avoid losing credits, and a stronger left frontal activation when encountering a CS indicting they had to chase a sheep to raise credits, findings suggestive of a role of frontal asymmetry (right vs left hemisphere activation) in avoidance or approach motivation. However, this finding that left frontal activation is stronger for approach conflicts with our results where Active Approach shows strong right frontal gamma. These contradicting results are likely due to differences in analysis methods (the use of relative alpha power) and can be further investigated in the future.

Alternatively, previous research has indicated that frontal beta-gamma (20–30 Hz) activity is thought to reflect a salience signal (i.e., motivational value signal) and has been associated with ongoing adjustments of behavior (HajiHosseini & Holroyd, 2015) and cognitive demand (Chen et al., 2012; Gilbert & Sigman, 2007; Lee et al., 2003). Further, studies have also shown frontal beta-gamma power increases are representative of increased attentional resources being applied to particularly important events (Bauer et al., 2014; Borghetti et al., 2021). In this context, our findings support the idea that frontal gamma reflects a manifestation of a motivational value signal that energizes behavioral adjustments when the CS are highly meaningful (i.e., must approach goal location to maximize rewards). Alternatively, the observed frontal gamma activity may also be related goal-planning for distal cues with high motivational value. For example, we recently demonstrated that beta-gamma power dissociated between uninformative cues (i.e. neutral feedback) and informative cues, and this lack of beta-gamma activity to neutral feedback coincided with a lack of trial-to-trial behavioral adjustment (P. Li et al., 2016). It is also worth noting that the early and late gamma response could highlight two distinct learning systems sensitive to avoidance behaviors (parietal gamma) and approach behaviors (frontal gamma). Although speculative, this reasoning predicts that the timing and topography of gamma activity should be correlated with measures of individual differences in threat and reward sensitivity and warrant future investigations.

## Conclusion

Despite the utility and simplicity of conventional fear conditioning paradigms, they nonetheless ignore the multitude of actions and decisions that animals and humans utilize in the real-world to avoid various threating situations. To our knowledge, our results demonstrate for the first time that the context of the CS differentially modulated theta and gamma activity and guided behavioral adaptation in the virtual RRI task. While the interpretation of these novel findings are admittedly speculative, these observations suggest that the neural oscillatory mechanisms underlying threat imminence can be investigated in humans and thereby a promising target for identifying and treating stress-related psychopathologies, specifically those characterized by maladaptive defensive behaviors. Such maladaptive processes have been identified across a spectrum of symptoms including but not limited to psychiatric disorders such as posttraumatic stress disorder (Maeng & Milad, 2017) and clinical anxiety (Suarez-Jimenez et al., 2021), as well as in high trait anxiety (Fung et al., 2019), childhood maltreatment (Lee & Hoaken, 2007) and suicidal patients (Millner et al., 2019). As such, identifying biomarkers of threat imminence could allow for therapeutic benefits, including objective measures of risk, diagnosis, and monitoring efficacy of treatment.

## Limitations

Although this research presents some of the first data using EEG and desktop virtual reality to examine oscillatory and behavioral dynamics during threat immense, future research may address some of the study’s limitations. First, although our sample was comparable to other EEG studies, a subset of participants did not learn the optimal strategy in both the threat and reward-CS conditions. While this pattern of results suggests that the RRI task is capable of uncovering individual differences in threat sensitivity, or possibly a general bias towards active responding, we hope to investigate this idea further using a larger sample of subjects as well as clinical and personality assessments. Second, traversing through any rendered environment via button presses or a joystick while physically immobile can clearly influence the degree of immersion and presence in the virtual environment during threat, and thus may not entirely resemble a “real-life” experience (Andreatta & Pauli, 2021). Several technological and methodological advances in electrophysiological research (mobile-EEG) and fully immersive virtual-reality (head mount display) have been made amenable for investigation in humans, and has started to see use in fear-conditioning paradigms (Stolz et al., 2019). We hope to leverage this advancement to investigate oscillatory dynamics in humans freely navigating a virtual environment under different threat imminence conditions.

## Footnotes

1 To note, the animal version of the RRI task was originally designed for the purpose of overcoming the limitations of current animal conditioning paradigms. Kyriazi and colleagues (2018) argued that current conditioning paradigms typically limit animal behavior to a single conditioned response (e.g. freezing), which can give the impression that CS automatically trigger conditioned responses when in fact it is the only option the experimental conditions allow. Since conditioning increases the likelihood that CSs will elicit a particular conditioned response, it is difficult to disentangle whether training-induced alterations in neural activity are related to the valence or sensory properties of the CS, the behavior they elicit, or a mixture thereof.

2 When considering Action as a factor in this analysis, a main effect of Action, F_(1,23)_=10.06 p<.005, η_p_^2^=0.30, and an interaction between Valence and Action, F_(1,23)_=5.56 p<.05, η_p_^2^=0.19, was observed. In contrast to the overall performance results presented above, the main effect indicated that participants were actually more accurate on trials requiring the optimal active response (x̄ =87%, SE=3) compared to the optimal passive response (x̄ =71%, SE=6; p<.005). Further, the interaction indicated that participants were less accurate at engaging the optimal passive response (x̄ =69%, SE=6) on avoidance trials in comparison to and active response (x̄ =88%, SE=2; p<.001). By contrast, for approach trials no differences were observed between the optimal passive (x̄ =72%, SE=7) and active responses (x̄ =83%, SE=4; p>.05). This interaction is likely due to passive behaviors being reinforced in the approach condition, as rewards were withheld if a response was detected. By contrast, passive behaviors were not reinforced in the avoidance conditions as participants simply needed to avoid the CS location to prevent the tone, regardless of response type.

