## Supplements for "Oscillatory correlates of threat imminence during virtual navigation"

**Supplement S1 – Verbal Instructions of the Task**

Verbal Task Instructions: “For this task, you will be ‘placed’ inside a room. While in this room, a blue color will appear either on the wall or the floor. When you see that blue color, you will need to make a decision. You can: 1) Stay in your current location, in which case do not press any button. Or 2) move to a different location in the room, in which case press the button on the keyboard that corresponds to the location to which you wish to move. Button ‘1’ on the number pad will take you to the left section of the room. Button ‘2’ will move you to the center of the room. And button ‘3’ will take you to the right section of the room. Note that each button corresponds to a specific location in the room – it does not move you right or left.”

Researcher asks if everything is clear up to this point, allowing for questions.

“Your goal is to figure out which decision will get you a reward, and which decisions will let you avoid getting the punishment. You know you have received the reward when you see the ‘$$$’ symbol on the screen. When you see it, it means you have earned 5 cents. While 5 cents does not sound like a lot of money, it can add up over the course of the task. The amount of money you earn during the task will be given to you as a bonus at the end of the study. That means that in total, there are three outcomes: reward, punishment, or nothing. Through trial-and-error, hopefully you will figure out which decisions are the best decisions to make.”

Researcher once again clarifies if there are any questions.

**Supplement S2 – Non-Parametric Statistical Methods**

Non-parametric permutation tests were run on the raw normalized power, based off the methods used by Cavanagh, Zambrano-Vazquez, & Allen (2012) using custom-made MATLAB scripts. First the clusters were identified, then multiple comparisons correction via permutation tests were run on the relevant cluster. To identify a cluster, paired-sample t-tests were computed at each time-frequency point (pixel) between active avoidance and passive avoidance conditions. For evoked Theta at P8 and total gamma at Pz, only pixels that survived the *p* < .01 threshold and had at least two adjacent pixels also *p* < .01 were retained (minimum cluster size = 3 frequency pixels by 3 time pixels). All adjacent significant pixels were then used to define a cluster. For total gamma at channel Fp2, pixels were thresholded at p<0.05 as no clusters were identified when using the criteria p<0.01.

Multiple comparison correction was completed using permutation tests of cluster-based thresholding, also referred to as “exceedance mass” (Nichols & Holmes, [2002](https://onlinelibrary-wiley-com.proxy.libraries.rutgers.edu/doi/10.1111/j.1469-8986.2011.01293.x#psyp1293-bib-0074)). The sum of all t-values in the pre-identified cluster were taken and used as the test statistic. To determine the p-value of the test statistic, permutations were used to create a null distribution of cluster summed t-values. For each permutation, t-tests were computed for each pixel in the cluster after 12 randomly chosen participants (half the participants) had their data switched between conditions. This was done under the null hypothesis that data between conditions are interchangeable. Data was switched within subject to account for the repeated-measures aspect of the study design. Once the null t-value of each pixel was calculated, the sum of all t-values in the cluster were taken to create a null distribution cluster value. One thousand permutations were run for each cluster to create the null distribution. The top and bottom 2.5% of the null distribution was used as a threshold providing a two-tailed 5% alpha level of family-wise error control for multiple comparisons. Supplementary figures 1,2,3 visualize the results.

|  | Evoked Theta | | | Total Gamma | | |
| --- | --- | --- | --- | --- | --- | --- |
| Condition | Peak latency  (msec) | Channel | Normalized Power | Peak latency  (msec) | Channel | Normalized Power |
| Active Avoidance | 245 | P4 | 0.76 | 336 | O1 | 0.08 |
| Passive Avoidance | 236 | P8 | 0.24 | 208 | Pz | 0.40 |
| Active Approach | 234 | P8 | 0.20 | 294 | Fp2 | 0.43 |
| Passive Approach | 234 | P8 | 0.44 | 316 | FC5 | 0.11 |
| All Conditions | 236 | P8 |  | 299 | Fp2 |  |

**Table S1.** Peak detection results on the normalized (non-log transformed) power. For each frequency and condition, the peak detection was run on data averaged across subjects and blocks. Peak results for “All Conditions” was run on data that was first averaged across all conditions (on top of block and subject). The search for peak power was restrained between 0-400msec after cue onset.

**Supplementary Figures**

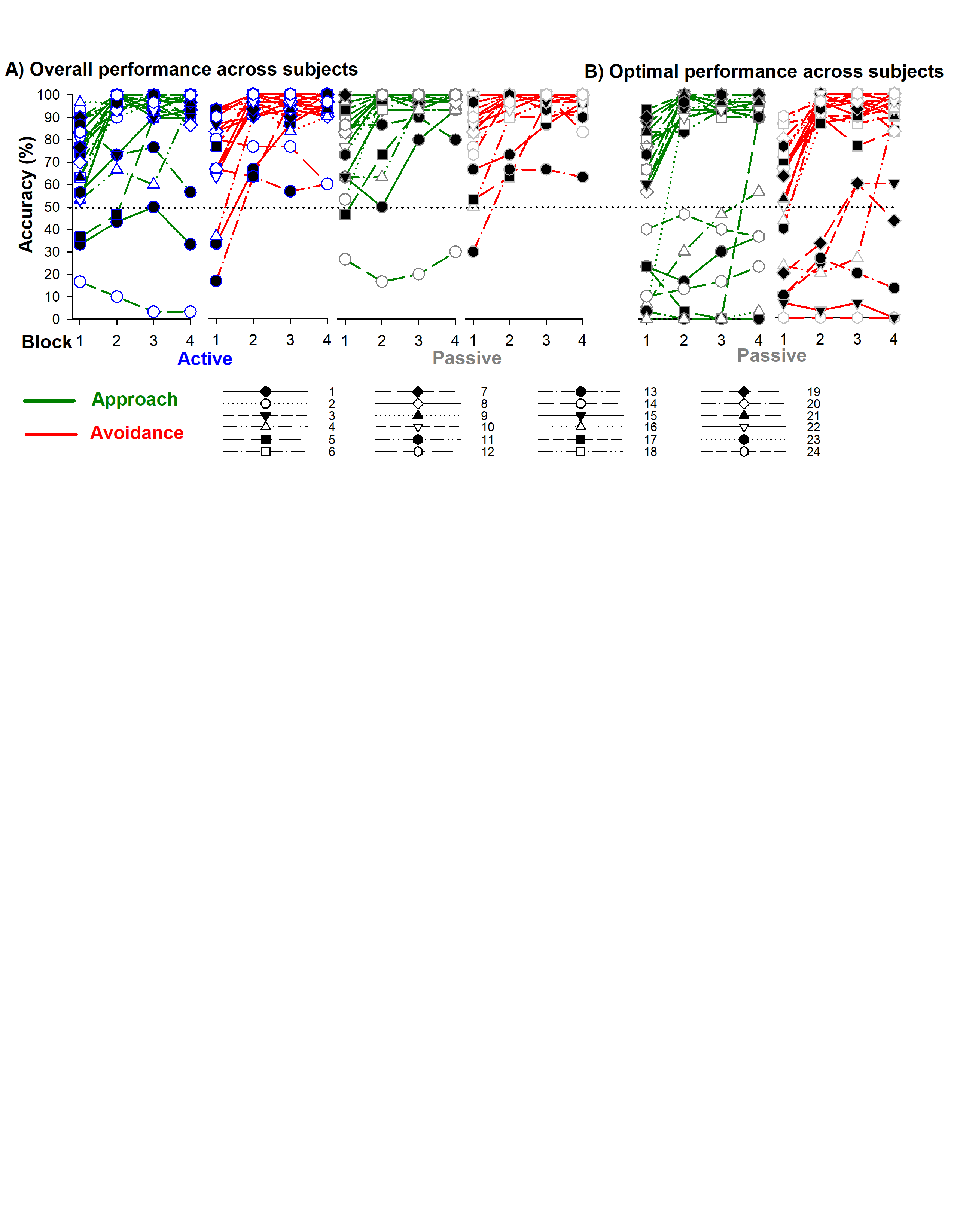

**Figure S1. Individual differences in RRI performance** **A.** Overall Active (Blue Symbols: left panels) and Passive (grey symbols: right panels) response accuracy across blocks for each subject as measured by percentage of Approach trials (green lines) receiving the monetary reward, and Avoidance trials (red lines) without receiving the aversive tone. **B**. Percentage of optimal passive (no button press) response in the approach (left panel) and avoidance (right panel) condition across blocks for each subject (n = 24).

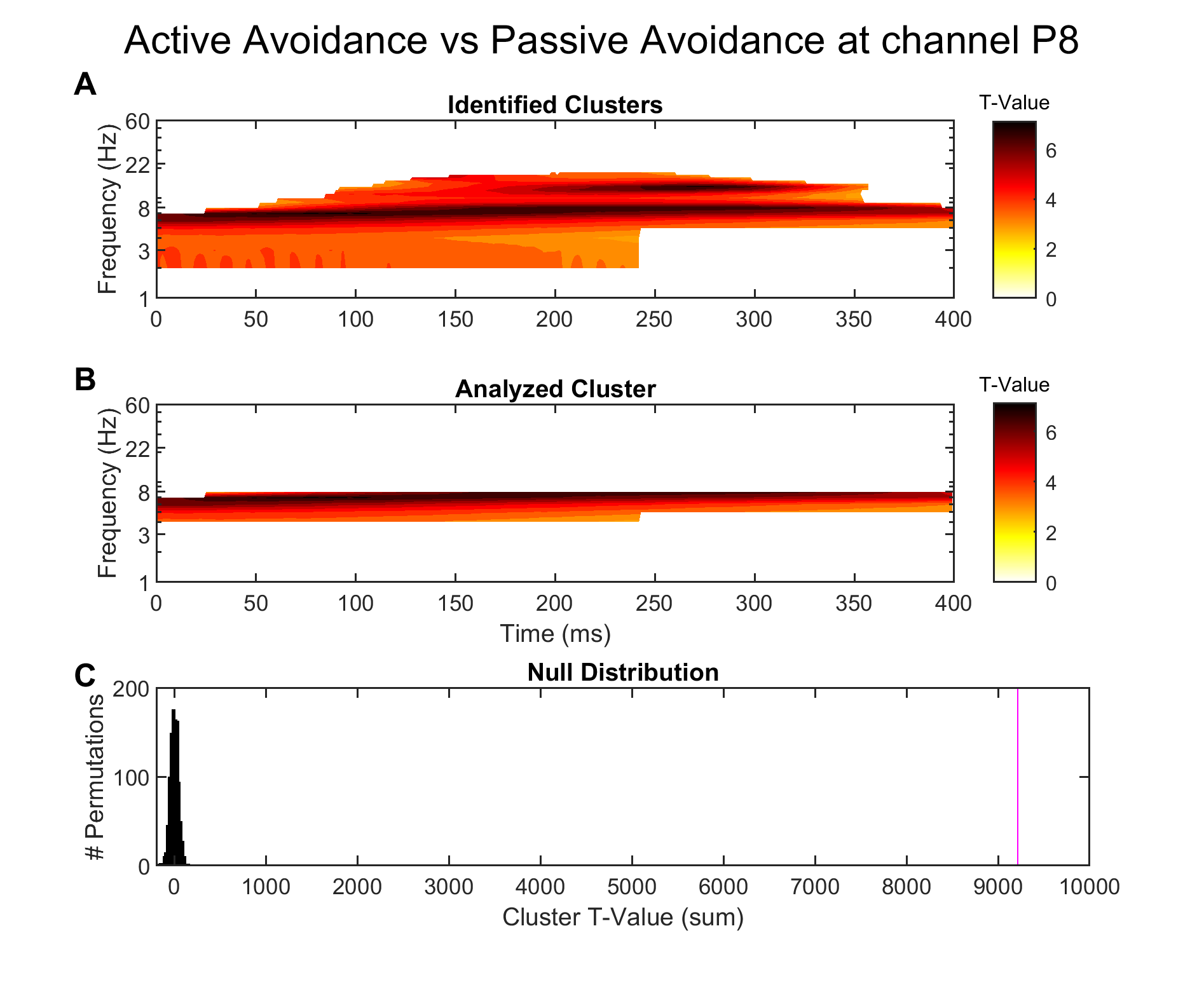
 **Figure S2. Cluster permutation results for evoked theta at channel P8 between Active Avoidance and Passive Avoidance conditions.** **A.** Cluster identified using the methods described in Sup. S2. **B**. Within the identified cluster, only pixels between the frequencies 4-8 Hz (i.e. Theta) were used for the permutation test to avoid inflating the summed T-value of the cluster. **C**. Histogram of the null T-value distribution, with the magenta line indicating the test statistic. The test statistic was significant (two-tailed, p = 0).

This permutation analysis supported the findings from the main text (Figure 8A). Cluster identification confirmed a significant difference in power that encompassed the theta range. This identified cluster spanned the entire search epoch (0-400msec), indicating that a real theta effect can be found somewhere within the cluster. Since the 211-261msec window of time used for the parametric statistics falls within that cluster, we can be more confident the window used during parametric statistics encompasses a real effect.

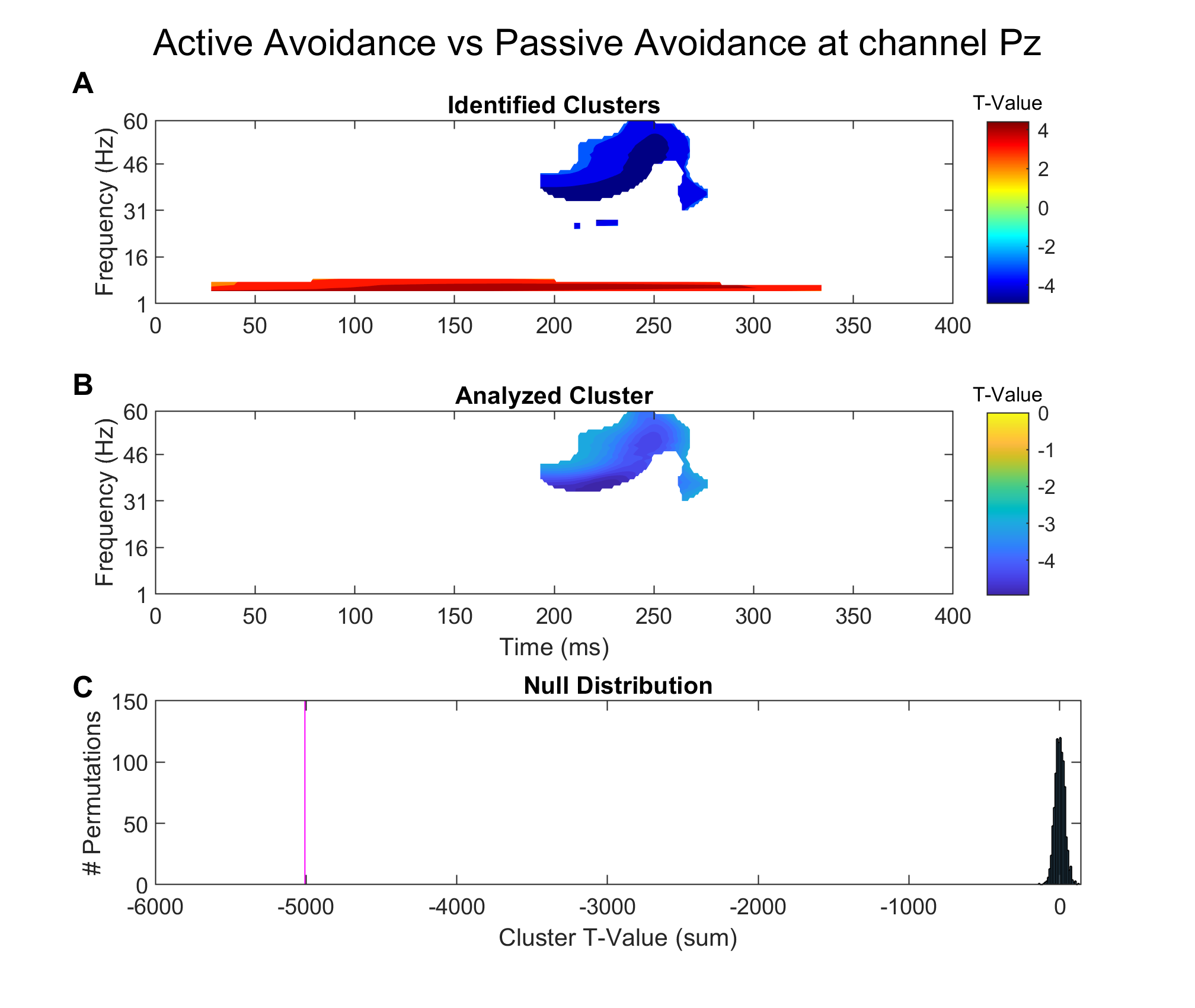

**Figure S3. Cluster permutation results for total gamma power at channel Pz between Active Avoidance and Passive Avoidance conditions. A.** Clusters identified using the methods described in Sup. S2. Multiple frequencies other than gamma were different between the two conditions **B**. Only the cluster encompassing the gamma frequencies (30-60Hz) were used for the permutation test **C**. Histogram of the null T-value distribution, with the magenta line indicating the test statistic. The test statistic was significant (two-tailed, p = 0).

This permutation analysis supported the findings from the main text (Figure 8B). Cluster identification confirmed a significant difference in power in the gamma range. This identified cluster spanned 192-277msec, indicating that a real gamma effect can be found somewhere within that range. Since the 183-233msec window of time used for the parametric statistics falls within that cluster, we can be more confident the window used during parametric statistics encompasses a real effect.

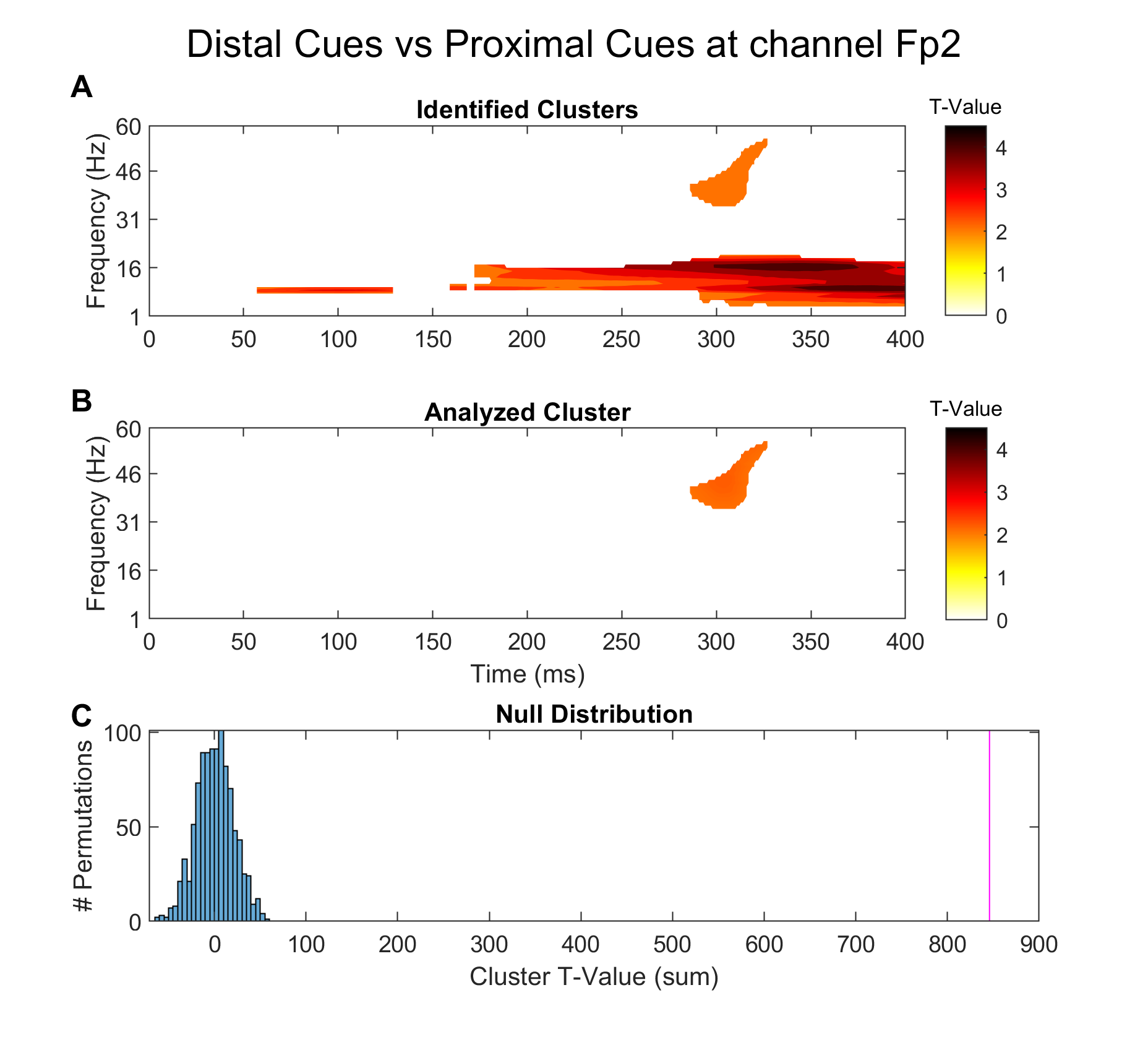

**Figure S4. Cluster permutation results for total gamma power at channel Fp2 between Distal Cues (Active Approach, Passive Avoidance) and Proximal Cues (Passive Approach, Active Avoidance)**. **A.** Clusters identified using the methods described in Sup. S2. Multiple frequencies other than gamma were different between the two conditions **B**. Only the cluster encompassing the gamma frequencies (30-60Hz) were used for the permutation test **C**. Histogram of the null T-value distribution, with the magenta line indicating the test statistic. The test statistic was significant (two-tailed, p = 0).

This permutation analysis supported the findings from the main text (Figure 8C). Cluster identification confirmed a significant difference in power in the gamma range. This identified cluster spanned 285-325msec, indicating that a real gamma effect can be found somewhere within that range. Since the 274-325msec window of time used for the parametric statistics falls within that cluster, we can be more confident the window used during parametric statistics encompasses a real effect.
